# Assessment of phytoplanktonic community and abiotic parameters of Rana stream in Mandi district, Himachal Pradesh, India

**DOI:** 10.1101/2023.12.07.570653

**Authors:** N. Bains, H.S. Banyal

## Abstract

This research paper gives insight into the physico-chemical characteristics of the Rana stream located in the Mandi district of Himachal Pradesh, India, with a particular emphasis on the diversity of diatoms. During the course of this study, eight genera of phytoplankton were identified. Statistical analysis using Pearson correlation unveiled significant associations between the physicochemical parameters and the different groups of diatoms. To assess the diversity of phytoplankton within the stream, several diversity indices were employed. Highest diversity was observed during the month of February. Furthermore, the calculation of water quality indices for the Rana stream yielded values falling within the range of 47.34 to 59.01. This range signifies that the water quality within the Rana stream can be categorized according to the Water Quality Index (WQI) scale, spanning from “good” to “poor.” It is worth noting that the presence of a diverse assembly of pollution-tolerant diatoms such as *Fragilaria, Navicula, Gomphonema*, and *Cymbella*, particularly during the month of March, which coincided with the period of the poorest water quality, is indicative of the eutrophic nature of the stream.

## Introduction

Streams represent pivotal inland freshwater reservoirs essential for addressing the escalating demand for water resources. They serve as a primary source of water for purposes including irrigation, potable water supply, and fisheries, contributing significantly to both economic and recreational value. Within the limnetic ecosystem, water quality is contingent upon a complex interplay of physical, chemical, and biological factors [1]. Phytoplankton species serve as vital indicators of the ecological condition of aquatic bodies, with their composition and dynamics playing a pivotal role in regulating biodiversity and energy flow within aquatic ecosystems. Phytoplankton, or microalgae, encompass phototrophic microorganisms characterized by uncomplicated nutritional requirements. They may be classified as either eukaryotes, such as green algae, or prokaryotes like cyanobacteria, as elucidated by [2].

The dynamics of phytoplankton exert a pronounced influence on trophic levels and the suitability of water for human utilization [3,4]. Phytoplankton form the fundamental link in the aquatic food chain, and nearly all dynamic attributes of aquatic bodies, including coloration, transparency, trophic state, zooplankton populations, and fish production, are substantially contingent upon the presence and behaviour of phytoplankton.

Diatoms constitute a prominent category within the realm of microscopic unicellular algae, and they rank among the most prevalent types of phytoplankton. Diatoms serve as crucial indicators of environmental modifications, where distinct species exhibit direct or indirect and highly responsive reactions to alterations in physical and chemical parameters. These parameters encompass nutrients, silicate, phosphorous, nitrogen, pH, light, and temperature [5]. Diatoms are extensively employed as bio-indicators in the evaluation of water quality due to their abbreviated generation times, and a multitude of species manifest specific sensitivities to ecological attributes [5,6]. Diatom communities have garnered significant favour as a tool for monitoring environmental conditions, both historical and contemporary, and are conventionally employed in investigations pertaining to the quality of aquatic ecosystems [7].

Limited information exists regarding the ecological patterns of phytoplankton and the state of water quality in the streams of Himachal Pradesh. Notably, the Rana stream, a significant tributary of the Beas riverine system, has received little scientific attention. Therefore, our research focuses on this specific stream.

The primary objective of this study was to assess the quality of water and its suitability for various purposes by utilizing various indices and comparing the results with established standards. Besides, diversity of phytoplankton especially diatoms is also considered.

## Area of study

The Rana stream is situated within the Jogindernagar region of the Mandi district in Himachal Pradesh, India. It is distinguished by its perennial nature, indicating that it maintains a continuous flow throughout the entire year. The origin of this stream is from Bir billing region located in Kangra district of Himachal Pradesh and the confluence point with Beas River is present at place Banaruawal of Mandi district which is approximately 27 km away from Jogindernagar region of Mandi district. This stream holds significance as a major tributary within the Beas riverine system.

**Fig. 1.**
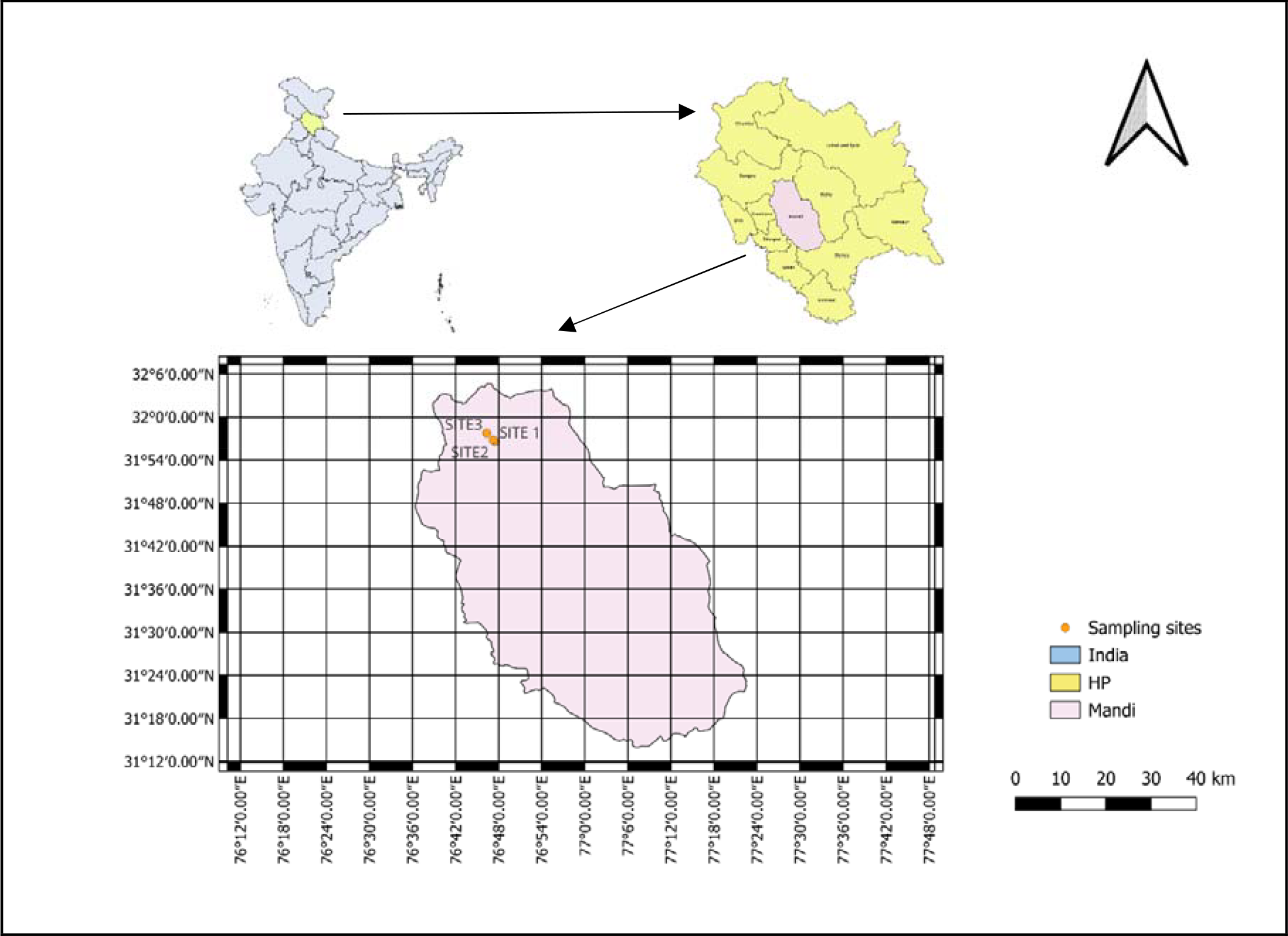
Study area of Rana stream.

## Materials and methodology

To evaluate the aqueous quality of Rana stream, monthly water specimens were procured from three designated sampling locations i.e. Near to Govt. Center Primaty school Khuddar, Near to Shiv temple, Banaun & Near to Sahib Bandgi Sant Ashram, Jogindernagar along the stream for a duration of 5 months, spanning from February 2023 to June 2023. Physicochemical parameters including water temperature, electrical conductivity (EC), total dissolved solids (TDS), dissolved oxygen (DO), and pH, were measured on-site using digital probes. The collected water samples were swiftly transported to a laboratory and temporarily stored for a brief period.

Within the laboratory, total alkalinity and total hardness were evaluated as per the methods prescribed by the American Public Health Association [8]. Plankton samples were collected using a plankton net made of No. 25 bolting silk and preserved them in a 4% formalin solution. Plankton density was calculated using a Sedgewick Rafter counting cell, and the identification of plankton species was carried out with standard reference books.

## Results & Discussions

In the course of the current study, a total of 8 genera of phytoplankton were identified. Among these, 5 genera (*Fragilaria, Navicula, Gomphonema, Cocconeis*, and *Cymbella*) belong to the class Bacillariophyceae, while the remaining 3 genera (*Spirogyra, Cladophora*, and *Ulothri*x) are classified under the class Chlorophyceae.

In the Rana stream, the pH levels reached their highest value in February (pH 9.9) and their lowest in June (pH 7.3), with an average value of 8.88 ± 0.92 (standard deviation). It is noteworthy that these pH values exceeded the permissible range (6.5–8.5) set by the Bureau of Indian Standards [9] and also greater than the values recorded by Gaury [10]. The elevated pH observed in February can be attributed to increased photosynthetic activity and the limited dilution capacity of the Beru stream during that month.

Significant positive correlations were observed between pH and Electrical Conductivity (EC) as well as Total Hardness (TH). pH also displayed non-significant positive correlations with Total Dissolved Solids (TDS), alkalinity, Dissolved Oxygen (DO), and chlorides. Conversely, a significant negative correlation was found between pH and water temperature. These findings are consistent with the work of Kumari & Kumar [11], who reported similar positive correlations between pH and alkalinity, as well as between pH and DO.

Furthermore, our analysis revealed that pH exhibited positive correlations with all the diatoms studied. Diatom *Cymbella* demonstrating a particularly significant relationship with pH. These results suggest that higher pH levels in the water are associated with larger populations of diatoms.

The reduced presence or complete absence of planktonic diatoms in the warmer areas of the lake is likely connected to the acidity of the lake water. Renberg & Hellberg [12], Flower & Battarbee [13], Charles [14], Huttunen & Thrkia [15], have suggested that the planktonic component is limited or non-existent at lower pH values. As Battarbee [16] has noted, planktonic diatoms tend to decrease in abundance when the pH of the water falls to around 5.5-5.8. The exact cause for this phenomenon remains uncertain, but one possible explanation could be a decrease in nutrient availability in acidic systems [17].

Alkalinity recorded its highest level in February, reaching 39.24 mg/L, while its lowest level occurred in June at 16.5 mg/L. The average alkalinity value during this period was 27.04 ± 7.25 mg/L. Notably, the recorded alkalinity values were lower than the standard value set by the World Health Organization [18].

Alkalinity exhibited a significant positive relationship with Dissolved Oxygen (D.O.) and a non-significant positive relationship with Total Hardness (TH) and chlorides. These correlations are in line with the findings reported by Jindal et al. [19], who also demonstrated a positive association between alkalinity and total hardness.

On the other hand, there was a non-significant negative correlation between alkalinity and water temperature, phosphates, and nitrates. These findings align with the work of Pandit et al. [20], who found a negative correlation of alkalinity with water temperature and nitrates, and Mustafa [21], who identified an inverse relationship of alkalinity with nitrates and phosphates.

Alkalinity exhibited a positive correlation with diatoms *Navicula, Fragilaria, Cymbella*, and *Cocconeis*. This correlation suggests that as the water becomes more alkaline, there is an increase in the presence of diatoms and the fish population. This observation aligns with previous research conducted by Hulyal & Kaliwal [22], who also noted that diatoms are typically more abundant in alkaline water.

Conversely, during present study it was found that alkalinity had a negative correlation with *Gomphonema*. Pedersen [23] observed that *Gomphonema* is indicative of low alkalinity, which means its abundance declines as alkalinity increases.

Temperature plays a critical role in various processes in aquatic environments, and in the Rana stream, the lowest water temperature was recorded in February (14.5°C), while the highest was observed in May (29.2°C), with an average value of 23.64 ± 5.33°C. The peak temperature in May can be attributed to increased sunlight intensity, reduced water levels, and clear atmospheric conditions.

During the present study, water temperature displayed a normal positive relationship with phosphates and nitrates, while it exhibited a normal inverse relationship with Dissolved Oxygen (D.O.) and chlorides.

Furthermore, water temperature showed a significant negative relationship with the diatom *Gomphonema* and a non-significant negative relationship with *Navicula, Cymbella*, and *Cocconeis*. This suggests that lower temperatures favor a higher abundance of diatoms and fish. These findings align with the observations made by Pandey & Verma [24] who noted an increased abundance of diatoms at lower temperatures.

The levels of dissolved oxygen (DO) in the Rana stream reached their lowest point in May at 6 mg/L, while they peaked in February at 8.9 mg/L, with an average value of 7.43 ± 0.97 mg/L. These DO values fall within the acceptable range established by the World Health Organization (WHO) of 4–10 mg/L [18].

D.O. exhibited a significant inverse relationship with phosphates and a non-significant inverse relationship with nitrates. These findings are consistent with the work of Pandit et al. [20] who reported a negative correlation between D.O. and nitrates, and Anshumali & Ramanathan [25], who observed an inverse relationship between D.O. and phosphate. In contrast, D.O. displayed a positive correlation with chlorides, which was also noted by Anshumali & Ramanathan [25].

D.O. exhibited a positive correlation with all diatoms. Dissolved oxygen plays a crucial role in supporting the growth of various flora and fauna. These findings align with the results observed by Jindal et al. [19], who reported a reduction in diatom abundance under conditions of low dissolved oxygen (DO).

Total Dissolved Solids (TDS) represent the cumulative concentration of dissolved salts in water. In the context of this study, the TDS level was found to be below the permissible limit of 1000 milligrams per liter (mg/L), with an average value of 71.50 ± 14.77 mg/L. The Rana stream displayed the highest recorded TDS value at 94 mg/L and the lowest at 56 mg/L.

The analysis revealed a significant positive correlation between TDS and alkalinity. This observation aligns with similar findings reported by Jindal et al. [26], which also documented a relationship between TDS and alkalinity. Conversely, TDS exhibited a significant negative correlation with phosphates, consistent with the results observed by Anshumali & Ramanathan [25].

In the study, the electrical conductivity (EC) levels in the Rana stream were carefully monitored. The highest EC value, recorded in March, reached 188 μS/cm, while the lowest, observed in June, was 108 μS/cm. The mean EC value during the investigation was determined to be 140.90 ± 30.83 μS/cm. Importantly, these EC values were well within the standard permissible limit of 2000 μS/cm, as reported by Kumar et al. [27].

During the investigation, several noteworthy correlations were observed. EC demonstrated a significant positive relationship with Total Dissolved Solids (TDS), alkalinity, and Dissolved Oxygen (D.O.). However, it exhibited non-significant correlations with Total Hardness (TH). The positive correlation with alkalinity aligns with the findings of Jindal et al. [19].

Conversely, EC displayed normal inverse relationships with phosphates, nitrates, and water temperature. These findings are consistent with the observations made by Anshumali & Ramanathan [25] regarding the relationships between EC, phosphates, and nitrates. Furthermore, Kumar et al. [27] also reported a negative correlation between EC and water temperature in Dal Lake.

Electrical conductivity (EC) displayed positive correlations with diatoms *Navicula, Gomphonema*, and *Cymbella*, while it exhibited negative correlations *with Fragilaria and Cocconeis*. This observation aligns with the findings of Bere [28], who noted the presence of *Gomphonema* in regions with higher EC. It was determined that EC is the most influential environmental factor affecting the distribution of diatom species, as reported by Pajuen et al. [29].

The highest total hardness (TH) was recorded in March at 62.67 milligrams per liter (mg/L), while the lowest was observed in June at 26.1 mg/L. The mean TH value over the study period was determined to be 48.28 ± 13.23 mg/L. It’s important to note that the total hardness levels in the stream remained well below the permissible limit established by Indian standards, which is 600 mg/L.

TH exhibited a significant negative relationship with water temperature, while it showed non-significant positive relationships with Dissolved Oxygen (D.O.) and chlorides. These findings are in line with the observations made by Kumar & Mahajan [30], who reported a positive association between TH and chlorides, and Pandit et al. [20], who observed a positive relationship between TH and D.O. TH showed a positive relationship with all the diatoms except *Fragilaria*.

In the Rana stream, the concentrations of chloride ions showed variations throughout the study. The lowest levels were recorded in June, measuring 16.24 milligrams per liter (mg/L), while the highest levels were observed in February, reaching 48 mg/L, with an average concentration of 22.55 ± 9.50 mg/L.

During the statistical analysis, it was determined that chlorides exhibited a non-significant negative correlation with phosphates and nitrates. This finding aligns with the observations made by Anshumali & Ramanathan [25]. Chlorides showed a significant inverse relation with Fragilaria while non-significant inverse relation with *Navicula, Gomphonema, Cymbella* and *Cocconeis*.

The concentration of phosphates in the Rana stream was quantified, resulting in a mean value of 0.28 ± 0.26 milligrams per liter (mg/L). The lowest phosphate concentration was observed in February, measuring 0.01 mg/L, while the highest concentration was recorded in June, reaching 0.65 mg/L.

Phosphates exhibited a non-significant positive correlation with nitrates. Furthermore, phosphates displayed positive correlations with diatoms *Navicula, Gomphonema, Cocconeis*, and *Fragilaria*, while showing a negative correlation with *Cymbella*.

Phosphate serves as a crucial nutrient essential for the proliferation and diversification of plankton within the aquatic environment, as reported by Namrata et al. [31]. Elevated phosphate concentrations accelerate the composition of diatoms, and phosphate nutrients play a pivotal role in facilitating cell division within diatoms. Katiyar et al. [32] observed that higher phosphorus concentrations lead to an increased rate of cell division in diatoms.

On the contrary, an excessive presence of phosphorus indicates a eutrophic environment, which can trigger harmful algal blooms, posing a lethal threat to aquatic life.

The concentrations of nitrates in the stream display seasonal variations, with the lowest levels observed in February at 0.01 mg/L and the highest levels in June at 0.33 mg/L, with an average concentration of 0.07 ± 0.09.

Nitrates exhibited a positive correlation with the diatom *Cocconeis* while showing a negative correlation with *Navicula, Fragilaria, Gomphonema*, and *Cymbella*.

Research by Atici and Obali [33] concluded that the pennate diatom *Cymbella* was tolerant to environmental factors Abubacker et al. [34] reported that species such as *Cymbella sp*. and *Gomphonema sp*. were tolerant to eutrophic conditions. Similar studies by Bhatt et al. [35], Jindal and Vatsal [36], and Ghavzan et al. [37] reported that forms like *Navicula cryptocephala* were pollution-tolerant and indicative of a high pollution load. During the present study, diatoms *Cymbella, Gomphonema*, and *Navicula* were also observed. Rott et al. [38] observed that *Fragilaria* is also highly resistant to pollution and can often thrive in eutrophic conditions.

### Diversity Indices

Diversity indices are valuable tools for assessing water pollution. Pristine waters are typically characterized by high species diversity, with a multitude of species sharing the ecosystem without one species dominating the population. Shruti et al. [39] concluded that pollution-induced stress tends to eliminate sensitive species while promoting the proliferation of tolerant species, leading to dominance.

Simpson’s Index (D) is a quantitative measure employed to assess species diversity based on dominance. This metric generates a numerical value ranging from 0 to 1, where 0 represents maximum diversity, and 1 signifies the complete absence of diversity. In simpler terms, a higher D value indicates lower diversity. In our ongoing investigation, we recorded the highest Simpson’s Index value for diatoms as 0.541 in May, while the lowest value was 0.415 observed in February.

Simpson’s Index of Diversity (1-D) calculates the probability that two randomly selected individuals from a sample belong to different species. This index also varies between 0 and 1, but in this context, a higher value reflects greater diversity. For diatoms, the maximum value of Simpson’s Index of Diversity (0.585) was observed in February, and the minimum value (0.459) was documented in May.

Much like Simpson’s Index, the Shannon Index of Diversity (H’) takes into account both species richness and evenness. An increasing value of the Shannon Index indicates a rise in both richness and evenness. For diatoms, the H’ values ranged from 0.714 to 1.053.

Sawan et al. [40] stated that values less than 1 suggest heavily polluted conditions, values between 1 and 3 indicate moderate pollution, and values greater than 3 point to clean water. Over the course of our study, the Shannon-Wiener index revealed that the water quality in the Rana stream ranged from heavily polluted to moderately polluted. The highest H’ value was recorded in February, signifying good water quality in that month.

Pielou’s Index quantifies the evenness of a community with numerical values between 0 and A value of 0 represents no evenness, while 1 signifies maximum evenness. In the case of diatoms, the J’ values ranged from 0.679 to 1, with the highest value, ‘1,’ observed in February, indicating maximum evenness during that month.

These diversity indices collectively suggest that the highest diversity of diatoms was recorded in February while the lowest diversity was observed in May month. Lowest value of Simpson index of Dominance (D) was observed in February month and highest value was observed in May month. High value of dominance possibly due to deteriorating water quality. Similar observations were made by Person [41], who reported that as water quality declines, the total number of species decreases, and a single species or only tolerant species dominate the community.

### Water Quality Index

The Water Quality Index (WQI) serves as a quantitative tool for assessing the state of water quality. This index is utilized to provide a numerical representation of water quality, with values ranging from 0 to 25 signifying “excellent” water quality, 25 to 50 indicating “good” water quality, 51 to 75 representing “poor” water quality, 76 to 100 denoting “very poor” water quality, and values exceeding 100 suggesting that the water is “unsuitable” for drinking as reported by Etim et al. [42]. Consequently, 10 physico-chemical parameters, including H_2_O temperature, pH, Total Dissolved Solids (TDS), Electrical Conductivity (EC), Dissolved Oxygen (DO), alkalinity, total hardness, chlorides, nitrates, and phosphates, were taken into consideration when calculating the WQI (see Table 5).

**Table 1.**
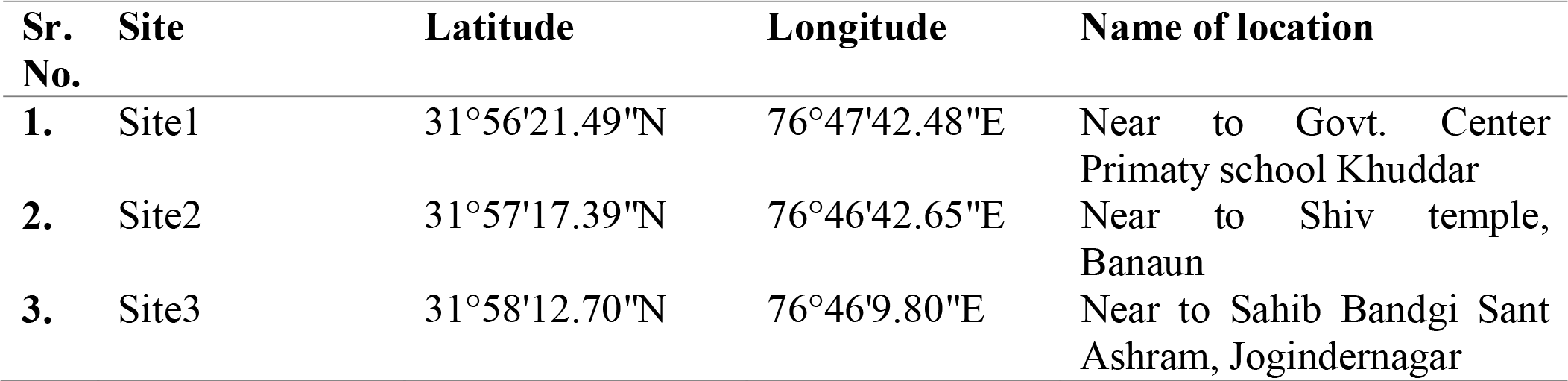
Location of sampling sites in Rana stream.

| Sr. No. | Site | Latitude | Longitude | Name of location |
| --- | --- | --- | --- | --- |
| 1. | Site1 | 31°56'21.49"N | 76°47'42.48"E | Near to Govt. Center<br>Primaty school Khuddar |
| 2. | Site2 | 31°57'17.39"N | 76°46'42.65"E | Near to Shiv temple,<br>Banaun |
| 3. | Site3 | 31°58'12.70"N | 76°46'9.80"E | Near to Sahib Bandgi Sant<br>Ashram, Jogindernagar |

**Table 2.** Physicochemical characteristics of Rana stream from (February 2023 to June 2023).

| Sr. No. | Parameters | Minimum | Maximum | Avg ± S.D. |
| --- | --- | --- | --- | --- |
| 1. | H<sub>2</sub>O Temp.(°C) | 14.5 | 29.2 | 23.64±5.33 |
| 2. | Ph | 7.3 | 9.9 | 8.88±0.92 |
| 3. | TDS (mg/L) | 56 | 94 | 71.5±14.77 |
| 4. | EC (μs/cm) | 108 | 188 | 140.9±30.83 |
| 5. | D.O. (mg/L) | 6 | 8.9 | 7.43±0.97 |
| 6. | Alkalinity (mg/L) | 16.5 | 39.24 | 27.04±7.25 |
| 7. | Total hardness (mg/L) | 26.1 | 62.67 | 48.28±13.23 |
| 8. | Chlorides (mg/L) | 16.24 | 48 | 22.55±9.50 |
| 9. | Phosphates(mg/L) | 0.01 | 0.65 | 0.28±0.26 |
| 10. | Nitrates (mg/L) | 0.01 | 0.33 | 0.07±0.09 |

**Table 3.** Exhibits the correlations that exist between the physicochemical characteristics of Rana stream.

|  | pH | EC | TDS | Alkalinity | TH | H<sub>2</sub>O Temp. | DO | Cl<sup>-</sup> | PO<sub>4</sub><sup>3-</sup> | NO<sub>3</sub><sup>-</sup> |
| --- | --- | --- | --- | --- | --- | --- | --- | --- | --- | --- |
| pH | 1 |  |  |  |  |  |  |  |  |  |
| EC | .650* | 1 |  |  |  |  |  |  |  |  |
| TDS | 0.601 | .992** | 1 |  |  |  |  |  |  |  |
| Alkalinity | 0.515 | .767** | .795** | 1 |  |  |  |  |  |  |
| TH | .891** | .636* | 0.586 | 0.63 | 1 |  |  |  |  |  |
| H<sub>2</sub>O Temp. | -.828** | -0.481 | -0.453 | -0.31 | -.684* | 1 |  |  |  |  |
| DO | 0.366 | .654* | .718* | .754* | 0.29 | -0.452 | 1 |  |  |  |
| Cl<sup>-</sup> | 0.091 | 0.353 | 0.349 | 0.236 | 0.22 | -0.123 | 0.281 | 1 |  |  |
| PO<sub>4</sub><sup>3-</sup> | -0.063 | -0.6 | -.648* | -0.627 | -0.152 | 0.006 | -.734* | -0.234 | 1 |  |
| NO<sub>3</sub><sup>-</sup> | -0.302 | -0.396 | -0.448 | -0.606 | -0.512 | 0.253 | -0.507 | -0.045 | 0.532 | 1 |

\* Correlation is significant at the 0.05 level (2-tailed).
\*\* Correlation is significant at the 0.01 level (2-tailed).

**Table 4.**
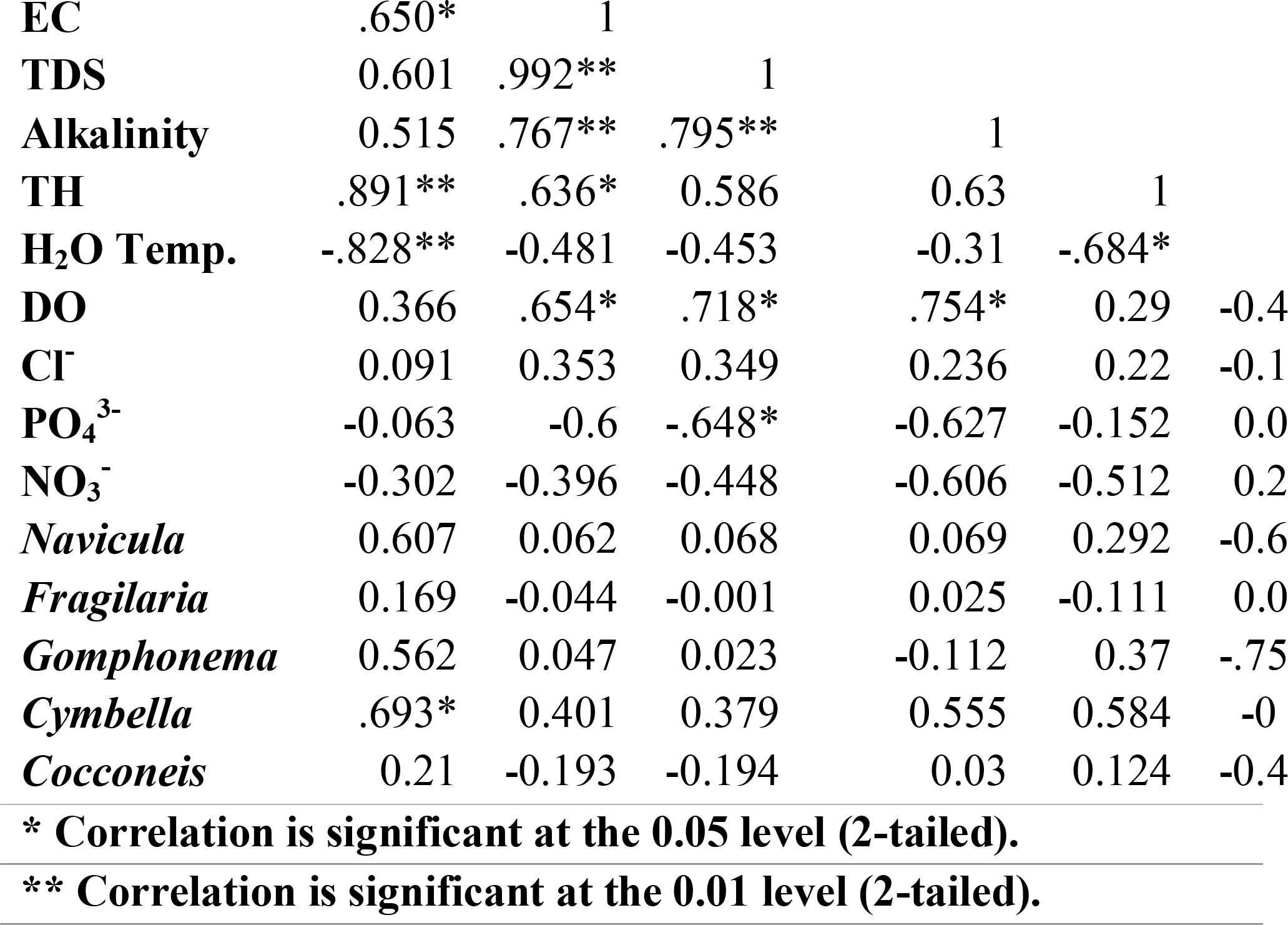
Exhibits the correlations that exist between the physicoch.

**Table 5.** Simpson’s Index (D), Simpson’s Index of Diversity (1-D), Shannon Index of Diversity (H’) and Pielou’s Index of Evenness of phytoplankton of Rana stream during the study period.

| INDEX NAME | MINIMUM | MAXIMUM |
| --- | --- | --- |
| Simpson's Index (D) | 0.415 | 0.541 |
| Simpson's Index of Diversity (1-D) | 0.459 | 0.585 |
| Shannon Index of Diversity (H') | 0.714 | 1.053 |
| Pielou's Index of Evenness | 0.679 | 1 |

**Table 6.** Water quality index of Rana stream.

|  |  |  | Feb. |  | March |  | April |  | May |  | June |  |
| --- | --- | --- | --- | --- | --- | --- | --- | --- | --- | --- | --- | --- |
| Parameters | S<sub>n</sub> | W<sub>i</sub> | Q<sub>i</sub> | W<sub>i</sub>Q<sub>i</sub> | Q<sub>i</sub> | W<sub>i</sub>Q<sub>i</sub> | Q<sub>i</sub> | W<sub>i</sub>Q<sub>i</sub> | Q<sub>i</sub> | W<sub>i</sub>Q<sub>i</sub> | Q<sub>i</sub> | W<sub>i</sub>Q<sub>i</sub> |
| **pH** | 8.5 | 0.12 | 111.18 | 12.80 | 113.53 | 13.08 | 111.76 | 12.87 | 94.71 | 10.91 | 91.18 | 10.50 |
| **TDS (mg/L)** | 1000 | 0.00 | 8.90 | 0.01 | 7.50 | 0.01 | 7.35 | 0.01 | 6.20 | 0.01 | 5.80 | 0.01 |
| **EC (ms/cm)** | 2000 | 0.00 | 8.83 | 0.00 | 7.50 | 0.00 | 7.35 | 0.00 | 5.88 | 0.00 | 5.68 | 0.00 |
| **D.O. (mg/L)** | 5 | 0.20 | 170 | 33.29 | 154 | 30.15 | 144 | 28.19 | 137 | 26.82 | 138.00 | 27.02 |
| **Alkalinity (mg/L)** | 120 | 0.01 | 28.60 | 0.23 | 24.86 | 0.20 | 23.51 | 0.19 | 20.29 | 0.17 | 15.42 | 0.13 |
| **Total hardness (mg/l)** | 300 | 0.00 | 18.21 | 0.06 | 20.89 | 0.07 | 18.06 | 0.06 | 13.77 | 0.04 | 9.55 | 0.03 |
| **Chlorides (mg/L)** | 250 | 0.00 | 6.69 | 0.03 | 13.86 | 0.05 | 7.95 | 0.03 | 7.04 | 0.03 | 9.56 | 0.04 |
| **Phosphates( mg/L)** | 1.5 | 0.65 | 1.40 | 0.91 | 23.67 | 15.45 | 24.43 | 15.95 | 19.00 | 12.40 | 24.67 | 16.10 |
| **Nitrates (mg/L)** | 50 | 0.02 | 0.02 | 0.00 | 0.09 | 0.00 | 0.11 | 0.00 | 0.04 | 0.00 | 0.42 | 0.01 |
| **WQI** |  |  |  | 47.34 |  | 59.01 |  | 57.31 |  | 50.38 |  | 53.83 |

(S<sub>n</sub>) Standards for drinking water (WHO, 2011).

The Water Quality Index (WQI) for the Rana stream falls within the range of 47.34 to 59.01, signifying that, according to the WQI scale, the water quality in the Rana stream can be classified as ranging from “good” to “poor.” Notably, the month of February exhibited the highest water quality, followed by the month of March, which was favored by the presence of a diverse diatom population. Surprisingly, despite the poorest water quality, the month of March displayed the highest diversity of diatoms, although it was lower than the diversity observed in February. This observation suggests that the current diatoms in the stream are tolerant to pollution and indicate the eutrophic condition of the water.

## Conclusion

The analysis presented here indicates that the water quality of the Rana stream varies from good to poor. When comparing the data from the Rana stream to international standards established by the World Health Organization (WHO), it becomes evident that certain physicochemical parameters, such as pH, Total Dissolved Solids (TDS), Electrical Conductivity (EC), alkalinity, Total Hardness (TH), chlorides, phosphates, and nitrates, are within the acceptable range for drinking water. However, parameters like Dissolved Oxygen (DO) exceed the permissible limits.

The water quality in the Rana stream was deemed good in the month of February, which can be attributed to the presence of a high diversity of diatoms. Subsequently, in the month of March, despite the water quality being poor, it supported the highest diversity of diatoms. This suggests that the diatoms found in the stream are tolerant to pollution and indicate a eutrophic condition of the water.

The computation of the Water Quality Index (WQI) for different months indicates that, without the implementation of appropriate measures for stream management, the water quality in the Rana stream is likely to deteriorate in the future. The presence of pollution-tolerant diatoms is indicative of the eutrophic state of the selected water body, which is primarily caused by an excessive influx of nutrients. Given the current state of the Rana stream, it is advisable to consider several conservation measures, including the development of a sewage management plan, the prohibition of fish-killing practices, and raising awareness among the local population.

